# Peaks and Distributions of White Matter Tract-related Strains in Bicycle Helmeted Impacts: Implication for Helmet Ranking and Optimization

**DOI:** 10.1101/2024.04.26.591318

**Authors:** Zhou Zhou, Madelen Fahlstedt, Xiaogai Li, Svein Kleiven

## Abstract

Traumatic brain injury (TBI) in cyclists is a growing public health problem, with helmets being the major protection gear. Finite element head models have been increasingly used to engineer safer helmets often by mitigating brain strain peaks. However, how different helmets alter the spatial distribution of brain strain remains largely unknown. Besides, existing research primarily used maximum principal strain (MPS) as the injury parameter, while white matter fiber tract-related strains, increasingly recognized as effective predictors for TBI, have rarely been used for helmet evaluation. To address these research gaps, we used an anatomically detailed head model with embedded fiber tracts to simulate fifty-one helmeted impacts, encompassing seventeen bicycle helmets under three impact locations. We assessed the helmet performance based on four tract-related strains characterizing the normal and shear strain oriented along and perpendicular to the fiber tract, as well as the prevalently used MPS. Our results showed that both the helmet model and impact location affected the strain peaks. Interestingly, we noted that helmets did not alter strain distribution, except for one helmet under one specific impact location. Moreover, our analyses revealed that helmet ranking outcome based on strain peaks was affected by the choice of injury metrics (Kendall’s tau coefficient: 0.58 ∼ 0.93). Significant correlations were noted between tract-related strains and angular motion-based injury metrics. This study provided new insights into computational brain biomechanics and highlighted the helmet ranking outcome was dependent on the choice of injury metrics. Our results also hinted that the performance of helmets could be augmented by mitigating the strain peak and optimizing the strain distribution with accounting the selective vulnerability of brain subregions, although more research was needed to develop region-specific injury criteria.

## Introduction

Traumatic brain injury (TBI) in cyclists is a concerning public health problem worldwide. In the United States, the annual number of emergency department visits for nonfatal bicycle-related TBI was around 596,972, with a large proportion of survived cyclists experiencing neurocognitive and behavioural sequelae [1]. In the Netherlands, the incidence of hospital admission due to bicycle-related TBI surged by 92% between 1998 and 2012, accompanied by an estimated annual economic cost of around 74.5 million Euros [2]. Injury data of cyclists in Sweden also showed that head injuries, in comparison to other trauma types, were more likely to cause permanent medical impairment [3]. Given the increased popularity of bicycling as a recreational activity and transportation mode [4], this TBI-related urgency in cyclists is expected to continuously grow and calls for effective prevention strategies [5].

Bicycle helmets are the most prevalent wearable gear for TBI prevention in cyclists by reducing the force acting on the head [6]. Existing bicycle helmets are engineered with the primary goal of passing the certification standard for commercialization purposes. For example, in Europe, the bicycle helmet is tested by dropping a helmeted headform onto a flat surface or kerbstone (i.e., direct impact) with the resultant peak linear acceleration (PLA) below 250 g (EN 1078). Similar standards exist in other countries, e.g., CPSC 16 CFR Part 1203 in the United States, GB 24429 in China, AS/NZS 2063 in Australia and New Zealand, etc. A detailed comparison of different bicycle helmet test standards is available in several recent reviews [7, 8]. However, existing helmet testing standards all focused on direct impacts with pass/fail criteria based on linear motion-related metrics (e.g., PLA, Head Injury Criterion (HIC)). Considering the prevalence of oblique impacts in real-life bicycle accidents [9] and the primary role of angular motion in brain injury mechanics [10, 11], several consortiums, such as the CEN Technical Committee 158 Working Group 11 in Europe, strived to improve the test standard by involving oblique impact scenarios (e.g., collisions against an angled surface) and angular motion-related metrics (e.g., peak angular velocity (PAV)) as the pass/fail threshold [12, 13]. Although multiple pilot bicycle helmet rating programmes with oblique impacts [14-21] have been proposed, none of them is officially involved in the testing standard yet.

Despite the progress in better reflecting the impact scenarios in bicycle accidents and brain injury mechanisms in the pilot rating programmes, one limitation is that the kinematic-based injury metrics (regardless of linear motion- or angular motion-based ones) are commonly used [15, 18-20]. Kinematic-based metrics only quantify the gross motion of the head and provide little information on how the brain and its subregions respond during helmeted impacts. To address this, researchers resort to finite element (FE) head models that can offer spatiotemporal detailed information on intracranial responses to external impacts of any magnitude and direction. Once gaining sufficient confidence of the model behaviour, the FE head model is an invaluable tool to assess the efficacy of different countermeasures based on localized mechanical loading (e.g., stress and strain) at the tissue level [22-30], complementing the kinematic-based injury assessment at the global level . A growing body of literature has evaluated the performance of bicycle helmets with the aid of the FE model [31-38]. For example, Fahlstedt, et al. [34] assessed the performance of one helmet in three accidental scenarios using the KTH Royal Institute of Technology head model and found that wearing a helmet reduced the maximum principal strain (MPS) in the brain by up to 45%. Abayazid, et al. [35] evaluated 27 bicycle helmets using another head model and found the performance of different protection technologies in terms of mitigating strain peaks was dependent on the impact direction. Several studies [14, 16, 17] proposed to integrate FE head models within the helmet rating programmes, although the capability of FE models for such a stringent application is at the forefront of the debate [39].

For brain responses during helmeted impacts, one fundamental question that is largely unknown is how different helmets alter the spatial distribution of tissue response. This is evidenced by the fact that the existing studies primarily focused on the peak value of tissue-based metrics, e.g., MPS [17, 29, 31, 34-38, 40, 41], maximum shear strain [32], maximal strain rate [35], maximal von-Mises strain [14], and maximal von-Mises stress [16]. Complementary efforts were noted to quantify the volume fraction of brain element with the strain peak over a certain threshold (i.e., the cumulative strain damage measure (CSDM) [42]) in helmeted impacts, but CSDM still could not provide location information of these “high-strained” elements. Even beyond the context of helmeted impacts, we found that only a handful of studies used FE models to investigate brain strain distribution using either synthetic angular loadings scaled from laboratory reconstructed impacts [43] or idealized sinus-shaped angular impacts [44]. Studying the brain distribution during impacts is motivated as the brain injury criteria might be region dependent. For example, one *in silico* investigation [27] reported that, out of 50 deep white matter (WM) regions of the human brain, the most vulnerable ones were genus of corpus callosum, cerebral peduncle, and uncinate fasciculus in football impacts. Several *in vitro* studies in rats found that the cortex were much less vulnerable than the hippocampus in terms of cell death [45, 46] and electrophysiological malfunction [47, 48]. Investigating the patterns of brain tissue responses in realistic helmeted impacts not only addresses an important research gap in brain biomechanics and may also provide new insights for helmet optimization.

Another important challenge when using the FE head model for helmet evaluation is how best to interpret the response of FE model. Today, no consensus has been reached on the optimal tissue-based metrics for brain trauma prediction (see the review by Ji, et al. [49]). The prevalence of concussions (an injury that often triggers axonal-related pathology [50, 51]) in cyclists [52, 53] motivated us to study the deformation of axonal fiber tracts in the assessment of bicycle helmets. Many experimental studies reported the stretch along the neuron/axon/fiber bundle instigated morphologic damage or functional alterations in the experimental tissue [54-56]. *In silico* studies from several independent groups also reported that the normal strain oriented along the fiber tract (i.e., tract-oriented normal strain, alternatively termed as tract-oriented strain, axonal strain, or fiber strain in the literature) demonstrated superior injury predictability than its counterparts (e.g., MPS) [22, 27, 57, 58]. Except for the tract-oriented normal strain, several *in vitro* studies suggested other types of tract-related strains as possible mechanical instigator of axonal pathology. For example, when imposing strain perpendicular to the neuronal direction, Braun, et al. [59] and Nakadate, et al. [60] noted tau pathology and axonal swelling, respectively, in the experimental tissue. LaPlaca, et al. [61] observed significant loss of neurites in the regime when the shear strain along the neuron peaked. In relevance to this multi-faceted injury mechanism, Zhou, et al. [62] presented a mathematical framework to comprehensively quantify the *in vivo* WM tract-related deformation during voluntary impacts based on three new tract-related strain measures, characterizing the normal strain perpendicular to the fiber tract (i.e., tract-perpendicular normal strain) and shear strains oriented along and perpendicular to the fiber tract (i.e., tract-oriented shear strain, tract-perpendicular shear strain, respectively). This framework could be extended to *in silico* simulations to employ tract-related strains as injury metrics in helmeted impacts. When relating to helmet evaluation, one interesting question that can be asked is how close the newly proposed tract-related strains discriminate the helmet performance to the prevalently used MPS.

The current study aimed to answer how different helmets influence brain strain patterns and how the helmet ranking outcome was affected by the choice of tissue-based injury metric. To this end, we extracted experimental kinematics from seventeen helmet models, each of which was tested at three impact locations. These impact loadings were imposed to an anatomically detailed FE head model with embedded axonal fiber tracts to predict the distribution and peak of four tract-related strains and MPS. The helmet performance was ranked based on five strain peaks. This study provided new insights into computational brain biomechanics and virtual ranking of helmet efficacy. It might also serve as a reference for helmet improvement, especially those intended to optimize brain strain distribution.

## Methods

### Laboratory helmet tests

All experiments were performed by the Folksam Insurance Group and described with greater detail available elsewhere [17]. In brief, seventeen commercially available bicycle helmets were purchased online or in-store from the Swedish market and tested in the laboratory to obtain impact kinematics. These seventeen helmets (Helmets A-Q, Fig 1A) were featured with different designs, of which eight (i.e., Helmets D, H, J, L, M, N, O, and Q) have rotational mitigation technologies used. During the test, the helmet was coupled with a 50^th^ percentile male Hybrid III headform with the chin strap fastened. The headform was in bare condition and its outer surface (i.e., a thick vinyl layer) was in contact with the helmet. The helmeted headform was dropped onto a 45° anvil with an impact speed of 6 m/s at three different impact locations (Fig 1B, referred to as XRot, YRot, and ZRot hereafter), each of which was expected to cause rotational motion primarily within one anatomical plane. The anvil was covered with a 40-grit sandpaper. The headform was instrumented with a 3-2-2-2 accelerometer package [63] to measure the linear and angular accelerations. The accelerometer samples were filtered with an SAE 180 filter [64] with a cut-off frequency of 1000 Hz. The impact kinematic data were expressed with reference to a headform-fixed coordinate system (shown in Fig 1B). In total, fifty-one helmeted impact experiments were performed. The recorded loading curves lasted for 30 ms, starting from the instant the helmet made contact with the angled surface.

**Fig. 1.**
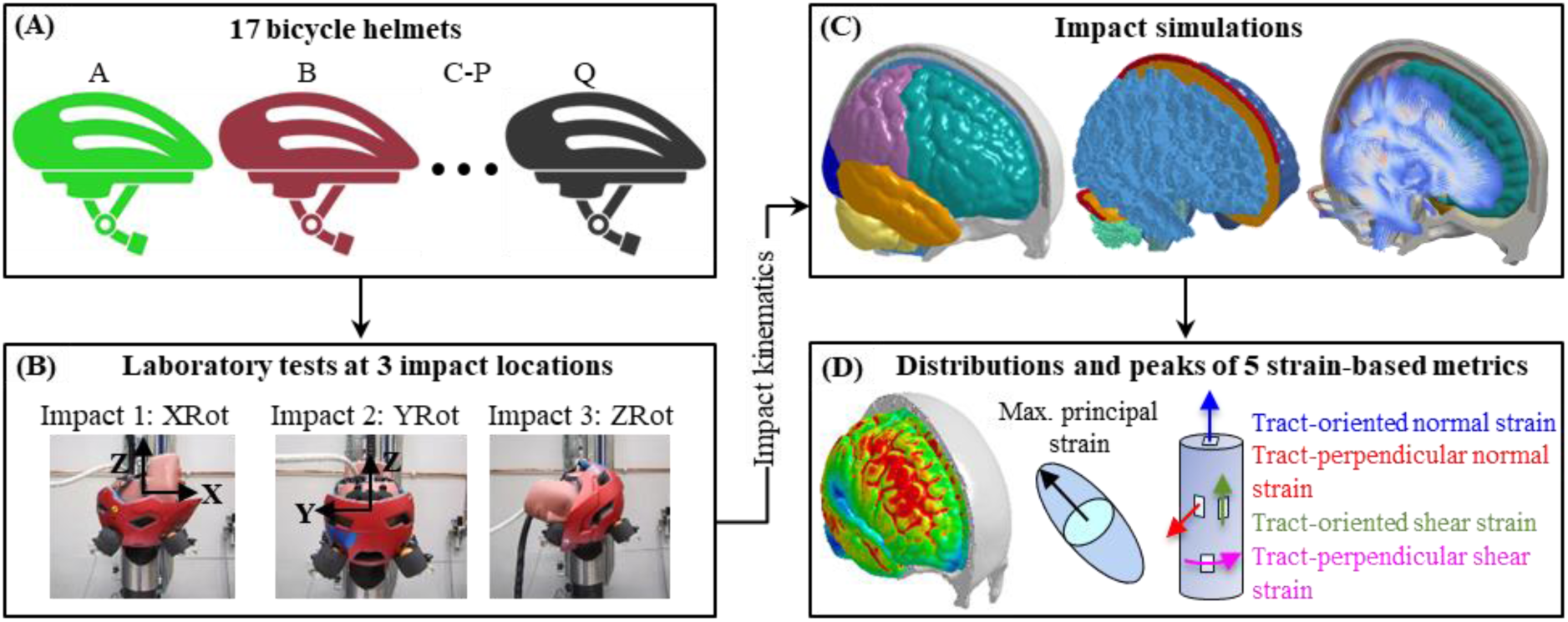
Illustration of study design and methodology. (**A**) Seventeen helmets (named as Helmet A to Helmet Q) available in the Swedish market were selected to be tested. (**B**) Three helmet impact locations were named as XRot for impact 1, YRot for impact 2, and ZRot for impact 3. The head kinematics were recorded by the instrumented accelerometers within the headform and expressed with reference to a coordinate system with the origin at the centre of gravity of the headform. (**C**) A computational head model with embedded white matter fiber tracts to simulate the helmeted impact secondary to the recorded impact kinematics. (**D**) Distribution and peaks of brain responses based on five strain-related metrics.

### Helmeted impact simulations

To estimate the localized brain strain responses, the experimental kinematics from the helmeted laboratory tests were imposed to an anatomically detailed computational head model (i.e., the ADAPT model in Fig 1C). The ADAPT model was developed previously at KTH Royal Institute of Technology in Stockholm using the LS-DYNA software with details about geometrical discretization, material modelling, interfacial representation, and model validation available elsewhere [65-67]. The model included gray matter, white matter (WM), cerebrospinal fluid, ventricles, skull, and meninges. To enable the calculation of tract-related deformation, the axonal fiber tract was embedded within the WM [65, 68]. Note that the fiber tract was only used in the post-processing procedure and had no mechanical contribution. Fifty-one impact simulations were performed, corresponding to the seventeen helmets tested in three impact locations. To simulate the helmeted impacts, the directional linear acceleration and angular velocity curves (shown in Fig A1 in Appendix) were imposed to one node located at the head’s centre of gravity and constrained to the rigidly modelled skull. All simulations lasted for 30 ms, each of which took 4 hours to solve using massively parallel processing version of LS-DYNA 13.0 version with 128 central processing units. The peak strain values of all elements were reached with the simulated impact duration.

To evaluate the brain responses in helmeted impact simulations, five strain-based metrics were employed (Fig 1D & Table 1A), including one brain tissue-level strain (i.e., MPS) and four WM tract-related strains (i.e., maximum tract-oriented normal strain (MTON), maximum tract-perpendicular normal strain (MTPN), maximum tract-oriented shear strain (MTOS), and maximum tract-perpendicular shear strain (MTPS)). The MPS was extracted for all brain elements (N = 513,198) and directly output from the simulation. The tract-related strains were only computed for WM elements (N = 213,321) with the exact mathematical equations presented in Appendix 1 in our previous study [62]. The peak values accumulated over time were extracted for all strain-based metrics.

**Table. 1.**
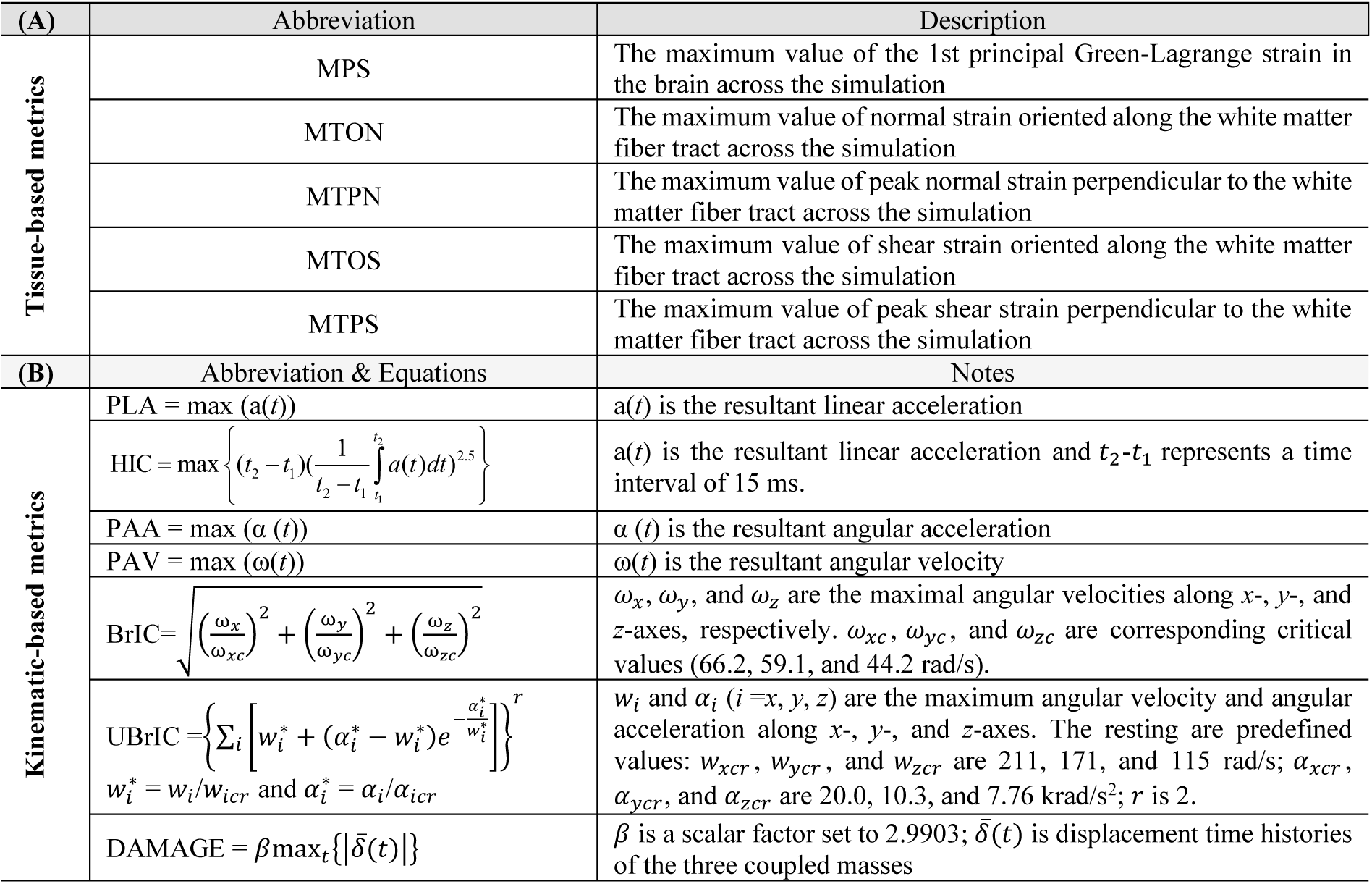
five strain-based metrics (**A**) and seven kinematic-based metrics (**B**) used in this study. Note that the coordinate system (*xyz*) used to compute the kinematic-based metrics is the same as the one in **Fig 1B**.

### Data analyses

For the impact kinematics in laboratory helmet tests, we evaluated the similarity of impact profiles (i.e., shape) between different helmets. Under a given impact location, Pearson’s correlations were conducted between the directional rotational velocity curves measured from any two helmets. For one impact, the time-history rotational velocity curves were expressed along X, Y, and Z axes and each curve lasted for 30 ms with a resolution of 0.05 ms, resulting in a sample size of 1,800 for each Pearson’s correlation. The current work tested seventeen helmets at three impact locations, leading to 408 Pearson’s correlation analyses (i.e., C_17_^2^ × 3 × 5).

We next quantified the peak responses of helmet impact loading based on seven kinematic-based metrics (Table 1B). These include two linear motion-based metrics (i.e., PLA and HIC [69]) and five angular motion-based metrics (i.e., peak angular acceleration (PAA), PAV [70], Brain Injury Criterion (BrIC) [70, 71], Universal Brain Injury Criterion (UBrIC) [72], and Diffuse Axonal, Multi-Axis, General Evaluation (DAMAGE) [73]). Equations for computing these metrics are summarized in Table 1B.

For the brain tissue responses in helmeted impact simulations, we quantified variations in peak strains among seventeen helmets by calculating the 95^th^ percentile value of MPS across the whole brain and four tract-related strain peaks across the WM for each impact simulation. The choice of 95^th^ percentile strain peaks was motivated to exclude the few elements with potential numerical instabilities, the same as the strategies in earlier works [74-76]. To statistically ascertain the influence of impact locations (XRot, YRot, and ZRot) on brain response, the peak strain results (non-normally distributed) were analysed with a Wilcoxon matched pairs signed rank test.

We ranked the seventeen helmets (Helmets A-Q) based on the strain peaks averaged across three impact locations, the same approach as adopted by Fahlstedt, et al. [39]. To assess the choice of strain-based metrics (five strains, Table 1A) on the helmet ranking outcome, Kendall’s tau coefficients were calculated based on the ranking results (10 different combinations). Pearson’s correlations were conducted to test the dependency between seven kinematic-based metrics and five strain-based metrics (35 tests), as well as among the five strain-based metrics themselves (10 tests).

To qualitatively visualize whether different helmets alter the strain distribution, we empirically identified the elements with the strain value ranked within the top 5% across the whole brain for MPS and the whole WM for tract-related strains for each simulation. We also normalized the element-wise strain by the 95^th^ percentile maximum strain result of the same simulation, similar to the approach of one early work [44]. Similar visualizations were illustrated for the corpus callosum to assess whether these seventeen helmet models altered the deformation pattern in this specific subregion that was often reported to be injured in TBI victims [77, 78] and frequently studied in biomechanical investigations [79-81].

To quantitatively assess the similarity of strain distributions in different helmeted impact simulations, Pearson’s correlation was conducted between element-wise strain peaks from the same impact location, akin to the method in one previous study [43]. This was done at the whole-brain level for MPS and the whole WM level for the four tract-related strains. The current work involved seventeen helmets, three impact locations and five strain-based metrics, resulting in a total of 2040 Pearson’s correlation analyses (i.e., C^2^_17_ × 3 × 5).

The threshold for significance was set to *p* < 0.05. The correlation level was determined by the coefficient value (*r*) (i.e., 0 ∼ 0.3 for weak correlation, 0.3 ∼ 0.7 for moderate correlation, and 0.7 ∼ 1 for strong correlation) [82].

## Results

### Helmeted impact kinematics

The impact pulses between different helmets were highly similar, except for one specific helmet at one impact location. For illustration purposes, we plotted the rotational velocity curves along three directions for Helmets A and Q tested at three impact locations in Fig 2A. In Impact 2 (i.e., YRot), both helmets had similar angular velocity profiles (*r* = 0.99) with the primary rotation along the Y-axis. This was similarly noted in Impact 3 (i.e., ZRot, *r* = 0.98), although the major rotation of both helmets was along the Z-axis. In Impact 1 (XRot), Helmet A primarily exhibited rotational velocity along the X-axis, whereas Helmet Q experienced significant rotational velocities across all three axes, resulting in an *r* value of 0.19. When expanding the correlation to all seventeen helmet models, we found that, under a given impact location, strong correlations (*r* > 0.7) in directional angular velocity curves were noted in all helmets (Fig 2B). We also noted exceptions in the pairs between Helmet Q and its 16 counterparts under XRot, of which all *r* values were less than 0.35. The detailed velocity curves for the fifty-one helmeted impacts were available in Fig A1 in the Appendix.

**Fig. 2.**
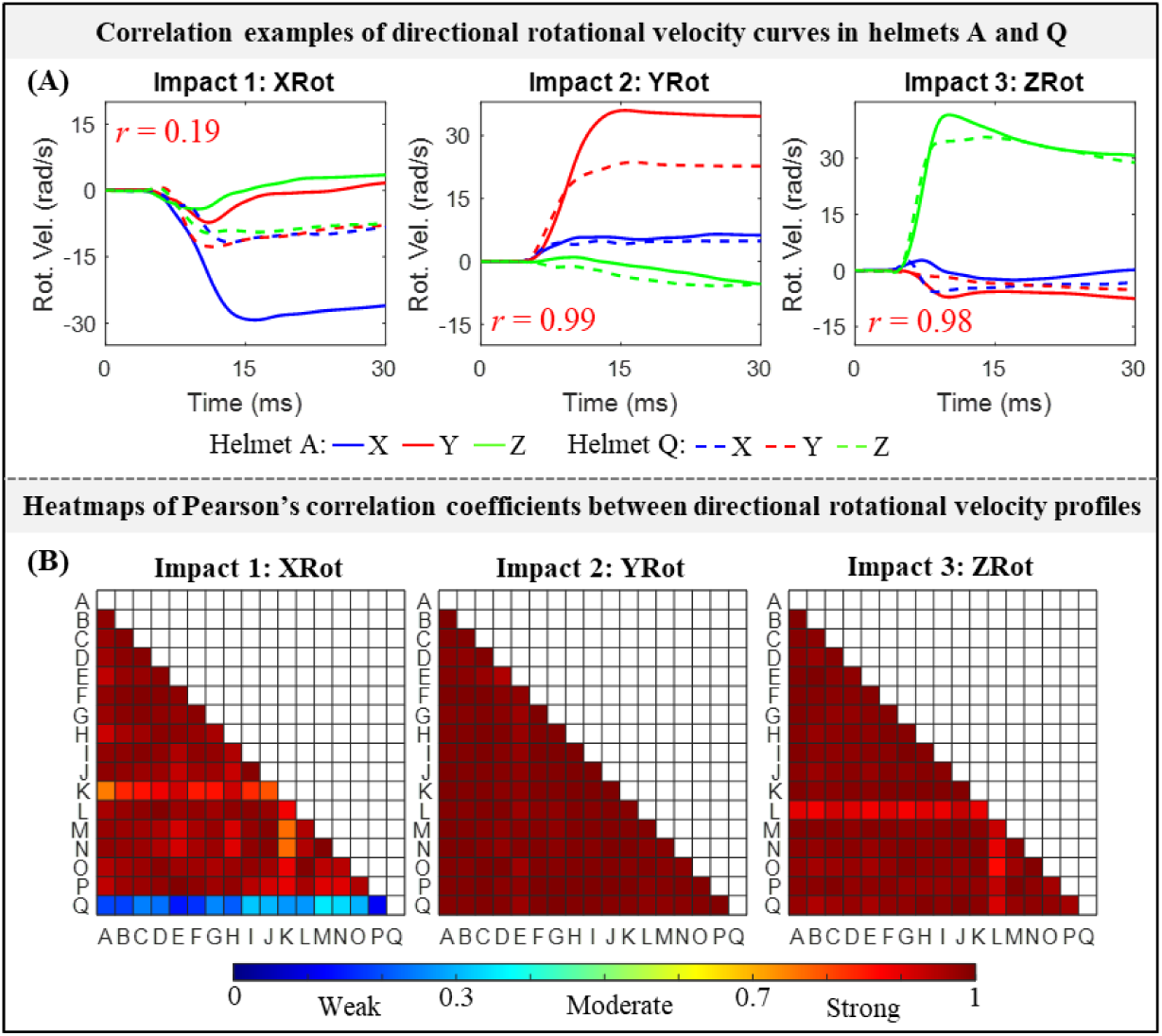
(**A**) Representative illustrations of directional rotational velocity curves in Helmets A and Q at three impact locations (i.e., XRot, YRot, and ZRot), and corresponding Pearson’s correlation coefficients (*r*). (**B**) Heatmap of Pearson’s correlation coefficient values for directional rotational velocity curves between seventeen helmeted impacts under the impact location of XRot, YRot and ZRot, respectively.

For the peak responses of helmeted impact loading, values of seven kinematics-based metrics were have been reported earlier [17, 39] and were not reiterated herein. To ensure the completeness of our study, we only presented a summary of seven kinematic-based metrics in Table 2.

**Table 2.**
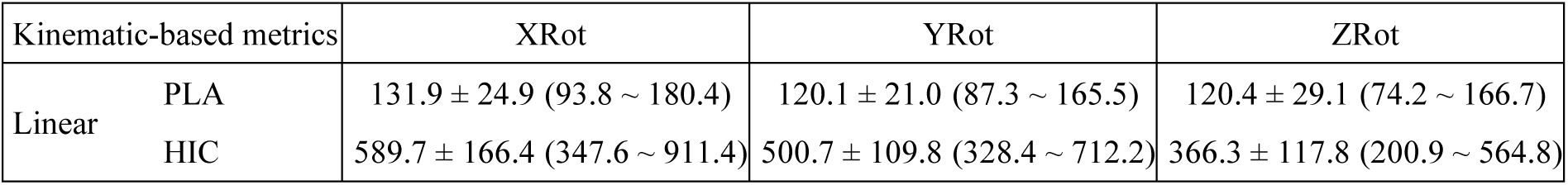
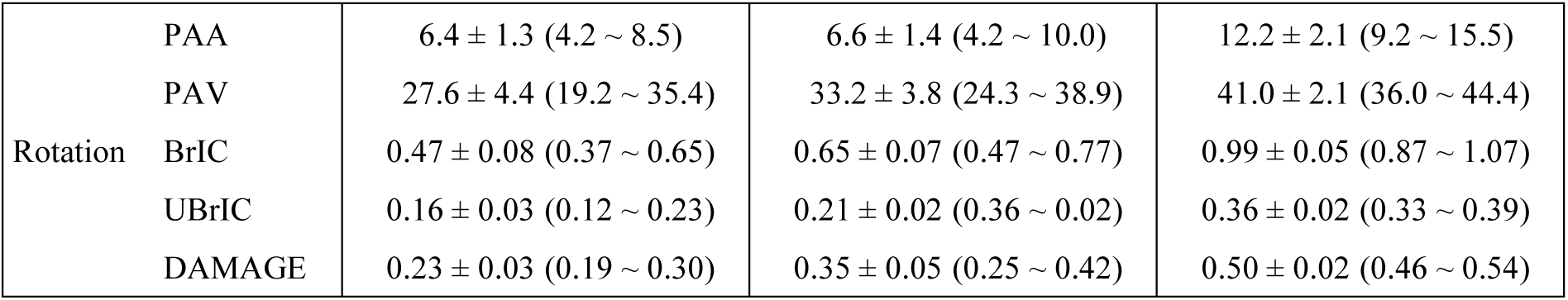
Summary of seven kinematic-based metrics in the form of mean ± standard deviation (range) for three impact locations (XRot, YRot, and ZRot).

### Brain strain peaks in helmeted impacts

The impact location affected brain strain peaks, but the effect was inconsistent among the five metrics (Fig 3A). In the seventeen helmets, strain peaks based on MPS, MTPN, and MTPS were significantly different (*p* < 0.001) in all three impact locations, in which the ZRot induced the highest values, followed by YRot and XRot. For MTON, the ZRot remained as the location with the highest value, while the YRot produced significantly smaller strain peaks than the XRot. When switching to MTOS, no significant difference (*p* = 0.19) was noted between XRot and YRot, while the ZRot instigated higher strain values than the other two impact locations. The exact values of 5 strain-based metrics for all 51 impact simulations were available in Table A1 in Appendix.

**Fig. 3.**
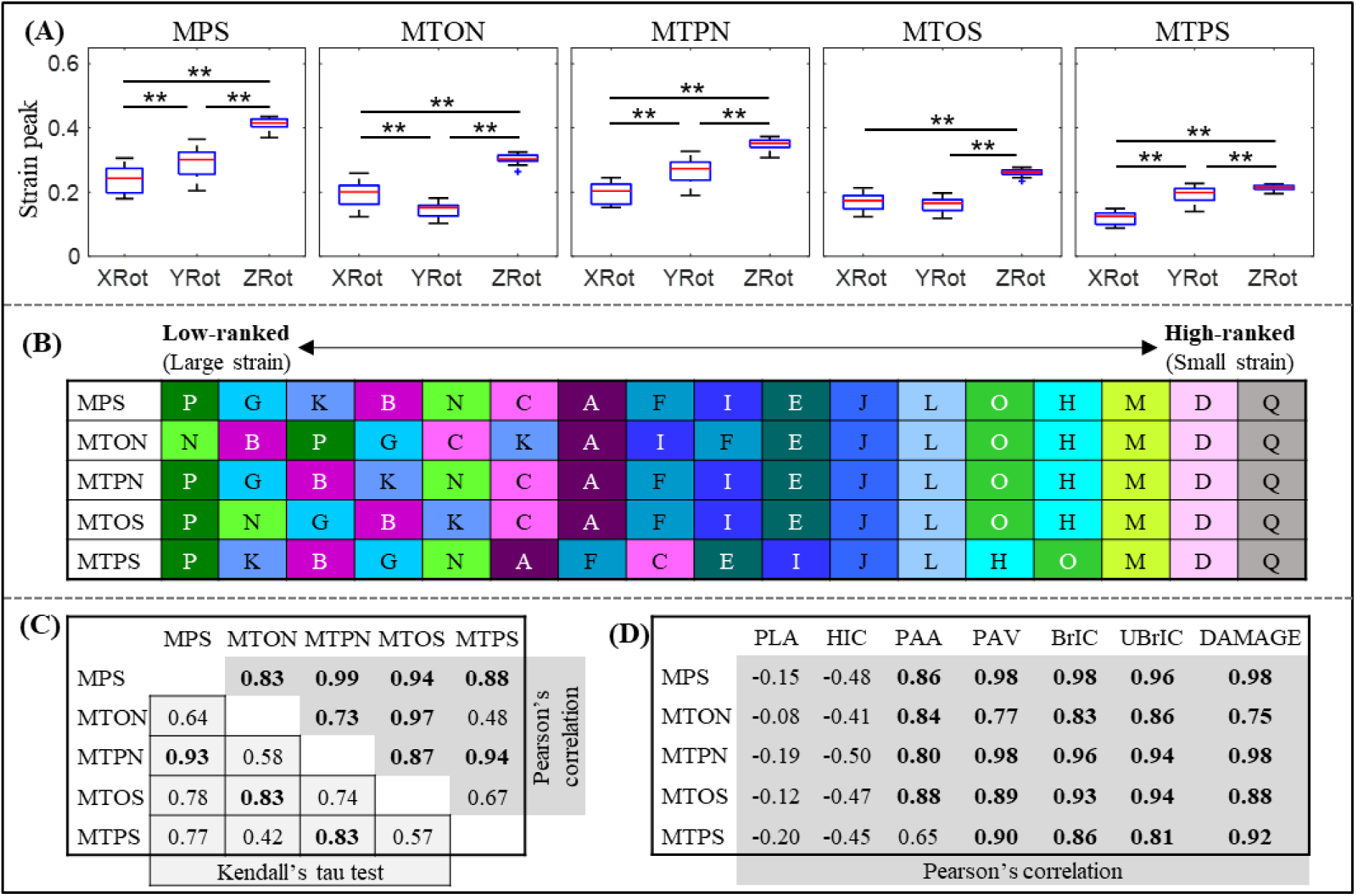
(**A**) Summary of five strain-based metrics in the form of boxplots for three loading locations (XRot, YRot, and ZRot) (\*\**p*<0.001, Wilcoxon matched pairs signed rank test). (**B**) Ranking of seventeen helmets (Helmets A-Q) based on strain peaks (i.e., 95^th^ percentile values averaged from three impact locations based on five strain types). (**C**) Below diagonal (cells colored in light grey with border lines in black): Kendall’s tau coefficient values to evaluate the sensitivity of helmet ranking on the choice of strain metrics with the coefficient values over 0.8 highlighted in bold. Above diagonal (cells in light gray with border lines unshown): Pearson’s correlation coefficient value (*r*) between five strain-related metrics. The coefficient values over 0.7 (i.e., strong correlation) were highlighted in bold. (**D**) Pearson’s correlation coefficient value (*r*) between five strain-related metrics and seven kinematic-based metrics. The coefficient values over 0.7 (i.e., strong correlation) were highlighted in bold.

For the helmet ranking based on the strain peaks averaged among the three impact locations, the same results were noted among the five strains for the high-ranked helmet, while significant disparities were noted in median-ranked and low-ranked helmets (Fig 3B). For example, Helmet Q was ranked the top (i.e., lowest strain), followed by Helmet D as the second and Helmet M as the third, independent of the strain type. For the low-ranked helmet, Helmet P was ranked lowest based on MPS, MTPN, MTOS, and MTPS, while Helmet N was ranked lowest based on MTON. We further quantified this ranking-related disparity using Kendall’s tau (Fig 3C, below diagonal) with the coefficient value varying from 0.42 to 0.93. Only three Kendall’s tau attained coefficient values over 0.8 (i.e., MPS vs. MTPN, MTON vs. MTOS, and MTPN vs. MTPS). For the interdependency among the five strains (Fig 3C, above diagonal), strong correlations (i.e., *r* > 0.7) were noted in all strain pairs, except for two cases (i.e., MTON vs. MTPS, MTOS vs. MTPS).

For the correlation between kinematic-based metrics and strain-based metrics (Fig 3D), all five strains exhibited strong correlations (i.e., *r* > 0.7) with all five angular motion-related kinematic metrics, except for one case (i.e., PAA vs. MTPS). For the linear motion-related kinematics, all five strains were either insignificantly (*p* > 0.05) or moderately/weakly (|*r*| < 0.6) correlated with PLA and HIC.

### Brain strain distributions in helmeted impacts

To evaluate whether the helmet altered the strain pattern, we identified the elements with the strain value ranked within the top 5% of the given simulation (termed as high-strain elements) and plotted the results of three representative helmets, i.e., Helmet Q (high-ranked), Helmet E (medium-ranked), and Helmet P (low-ranked) (Fig 4A). It can be noted that the distribution of high-strain elements depended on the impact location and strain type. Interestingly, the helmet model did not change the location of high-strain elements. Taking the MTON in YRot as an example, high-strain elements were consistently noted in the cerebral WM in the frontal and occipital lobes, and cerebellum WM for all three helmets. However, one exception was noted in XRot between Helmet Q and the other two helmets, regardless of the strain type. Using MTPN as an illustrative metric, high-strain elements were wholly located within the cerebral WM for Helmet Q, while partial high-strain elements were noted in cerebellum WM for Helmets E and P. While scrutinizing the normalized strain distribution (Fig 4B), we also noted similar results that under the same impact location, the helmet model did not affect the normalized strain pattern with the only exception noted in XRot between Helmet Q and the other two helmets. Similar illustrations for the corpus callosum were showed in Fig A2 in Appendix. Same as the finding at the whole brain or whole WM level, different helmets did not alter the strain pattern in the corpus callosum (see Fig A2 in Appendix), except for the Helmet Q at the XRot. Taken the MTPN as an example, high-strain elements were clustered at genu region for Helmet Q and distributed at the adjunction between the splenium and midbody for Helmets P and E (Fig A2A).

**Fig. 4.**
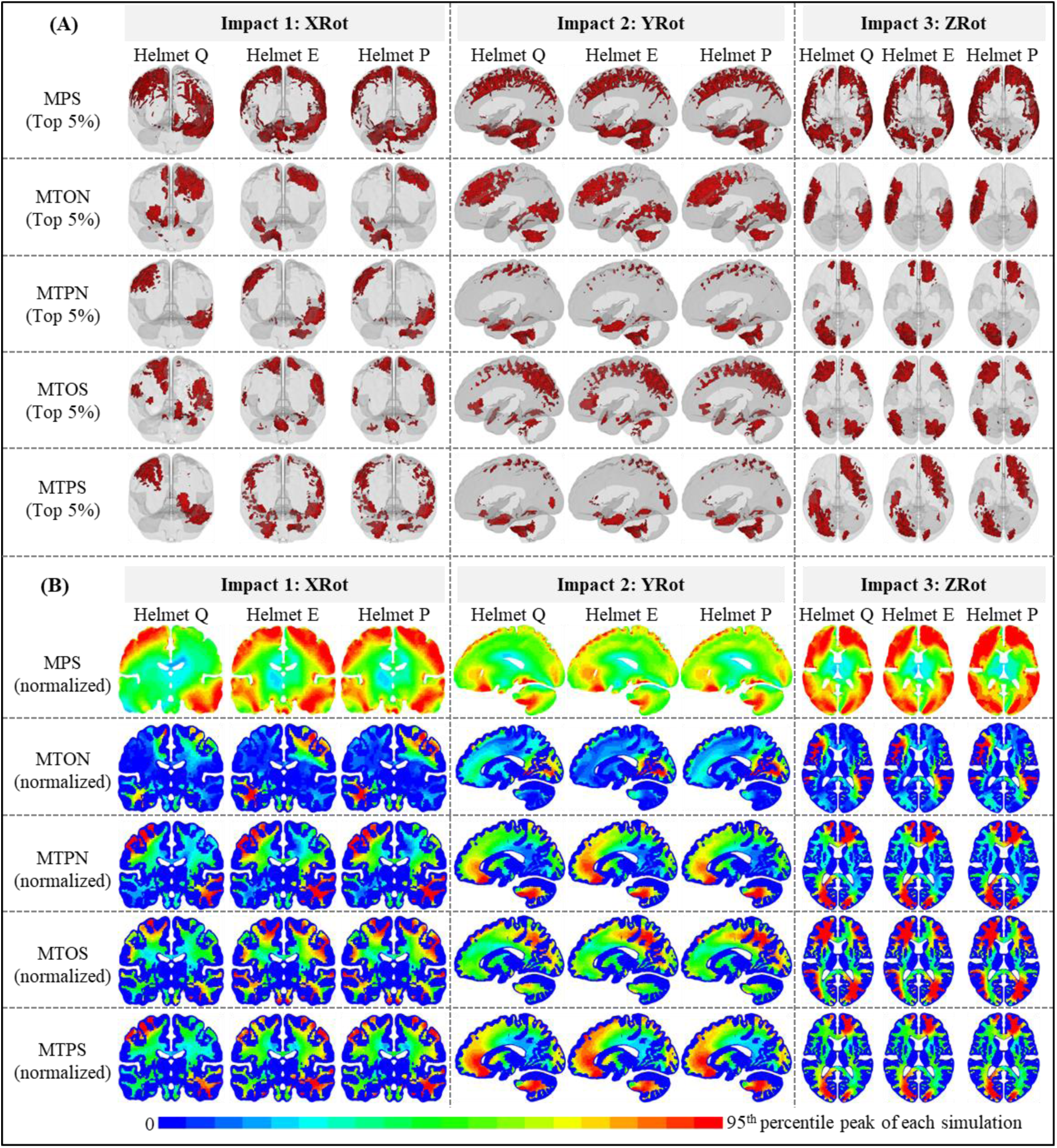
Brain strain distribution in nine representative impacts (i.e., Helmets Q, E, and P in XRot, YRot, and ZRot). (**A**) Brain elements with the MPS values within the top 5% and white mater elements with the tract-related strain values within the top 5%. To facilitate the visualization, the pia mater is shown in transparent. (**B**) Normalized brain strain contours in coronal, sagittal, and horizontal views for XRot, YRot, and ZRot, respectively. Note that, as indicated by the upper limit of the legend bar, the strain in each contour was normalized by the 95^th^ percentile strain values (available in Table A1 in Appendix) from the same simulation.

We further quantified the similarity in strain distribution by performing Pearson’s correlation in element-wise strain peaks. Representative plots for the correlations between Helmet D and Helmet E in XRot were shown in Fig 5A with *r* values as 0.95 for MPS, 0.98 for MTON, 0.96 for MTPN, 0.95 for MTOS, and 0.92 for MTPS. We next extended the correlation to all seventeen helmets with the *r* value plotted in the form of heatmap in Fig 5B for XRot, Fig 5C for YRot, and Fig 5D for ZRot. Under a given impact location, strong correlations (*r* > 0.7) in the element-wise strain results were noted in all seventeen helmets, regardless of the strain type. The only exception was noted between Helmet Q and the other 16 helmets, in which only weak to moderate correlations (*r* < 0.7) were noted. Taken together, the correlation results of element-wise strain peaks quantitatively verified that the helmet did not change the strain pattern except for Helmet Q in XRot.

**Fig. 5.**
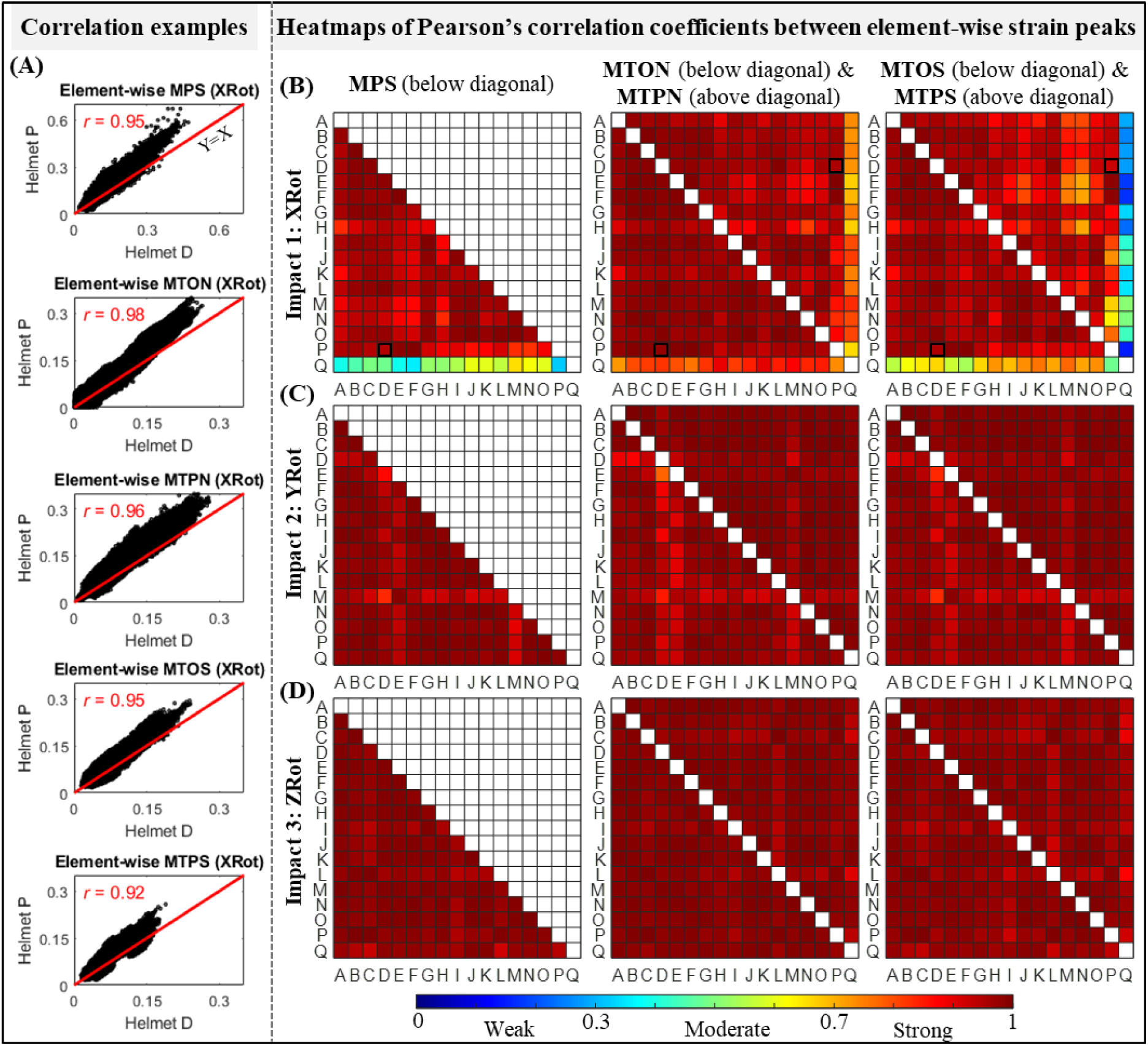
Pearson’s correlation coefficient (*r*) between element-wise MPS, MTON, MTPN, MTOS, and MTPS from 51 helmeted impacts (three impact scenarios, i.e., XRot, YRot, and ZRot, and 17 helmets, i.e., helmets A-Q). (**A**) Representative illustrations of Pearson’s correlation for element-wise MPS, MTON, MTPN, MTOS, and MTPS from impact simulations with kinematics obtained from helmets D and P in XRot. (**B-D**) Heatmap of Pearson’s correlation coefficient values for element-wise strain results between seventeen helmeted impacts under the location of XRot, YRot and ZRot, respectively. Note that in subfigure B, the cells corresponding to the five specific correlations in subfigure A are highlighted with border lines in bold.

## Discussions

The current study investigated the peaks and distributions of five strain-based metrics in fifty-one helmeted impact simulations (seventeen helmet models, three impact locations). Our results showed both the helmet model and impact location affected the strain peaks. The helmet ranking outcome based on the strain peaks was affected by the strain types. Interestingly, under a given impact location, the helmet did not change the strain distribution with only one exception noted. This study reinforced the importance of mitigating the strain peaks in helmeted impacts and hinted at a possible new direction of helmet improvement by optimizing the strain distribution with accounting the selective vulnerability of different brain regions. Our work also provided new insight that the helmet ranking outcome was dependent on the choice of injury metrics.

Our results on the similarity of brain strain patterns under the same impact location were generally agreed with the findings in previous *in vivo* imaging and computational studies. Knutsen, et al. [83] implemented tagged magnetic resonance imaging to measure MPS and MTON in twenty voluntary impacts with two head motions, i.e., ten neck rotations with primary angular motion within the axial plane and ten neck extensions with primary angular motion with the sagittal plane. It was found that strain patterns based on MPS and MTON were spatially consistent across volunteers for the same head motion. By excising one three-dimensional FE head model along the coronal plane by 44 synthetic angular velocity profiles with various shapes but the same PAV (23.4 rad/s) and duration (46.3 m/s), Zhao, et al. [43] found that the MPS pattern remained largely consistent with the MPS peaks significantly changed. Similar findings were also reported by Carlsen, et al. [44] using three planar FE head models secondary to 100 idealized sinus-shaped angular loadings along the sagittal, coronal, and axial planes, respectively. Our study extended previous efforts by using realistic loadings recorded from seventeen bicycle helmets in three distinct impact locations. These impact locations represented the most common collision sites in bicycle accidents [84-91]. Our results also further confirmed the early finding the strain peak was affected by the helmet model and impact locations. Relevant discussion on this has been extensively reported in the literature [31-38] and was not repeated here.

Our study provided important implications for helmet improvement. The large variations in MPS at the whole brain level and tract-related strain peaks at the whole WM level (Fig 3) implied that existing helmets could still be improved by mitigating the strain peak, as has been reported before [34-36]. This implication was based on the rationale that the peak strain represented the worst scenario and was most likely to cause injury. However, it did not recognize that different brain regions responded differently to identical mechanical stimuli. For example, Morrison III and colleagues stretched *in vitro* cultured hippocampal and cortical tissues from rats and found, compared to the cortex, the hippocampus was more susceptible to injury in the form of cell death [45, 46] and electrophysiological impairment [47, 48]. Even among the subfields of the hippocampus, cornu ammonis 1 and 3 were less resistant to cell death than dentate gyrus, when exposed to the identical strain [46]. One recent modelling study of the human brain by Zimmerman, et al. [92] also reported that distinct distribution of biomechanical loading during head impacts might be associated with different neurologic impairments, i.e., high strain and strain rate in brainstem might cause loss of consciousness, while large deformation in the motor cortex might instigate dystonic posturing. In light of this, we studied the brain strain distribution in fifty-one helmeted impacts, qualitatively (Fig 4) and quantitatively (Fig 5), with the involvement of all brain elements for MPS and all WM elements for four tract-related strain peaks. Our results found evident similarity in strain patterns in all seventeen helmets (with only one exception that was discussed later), indicating that different helmets did not change the path of mechanical force transmission within the brain during impacts [44]. Such results hinted that the protective performance of existing helmets might be able to be further augmented by optimizing strain distribution by accounting the selective vulnerability of brain subregions, i.e., redistributing the brain strain with a relatively large dose in the region that was more resistant to injury and a relatively small dose in the region that was more prone to injury. It should be clarified that such an improvement was at the conceptual stage as the exact values of region-specific injury criteria for the human brain were largely elusive today. Initial attempts have been noted to use human FE head models to derive region-specific injury risk functions [22-28, 58, 93, 94]. However, these risk functions were derived by correlating continuous strain-based metrics to binary-diagnosed injury outcomes (concussion or no-concussion), where definitive pathologies at specific regions were lacking. Severe under-sampling of non-injured cases was often noted while using FE head model to develop risk functions. As was critically reviewed by Ji, et al. [49] and Siegmund, et al. [95], this might lead to overestimated injury probability when using these risk functions to assess injury. Thereby, further evaluation was needed to assess the reliability of these aforementioned region-specific risk functions derived from human FE head models.

The lack of similarity in strain patterns between Helmet Q and the other sixteen helmets in XRot could be explained by the disparity in impact kinematics. As was quantified in Fig 2 and also plotted in Fig A1 in Appendix, Helmet Q had comparable angular motion in all three anatomical directions with the peak angular velocity as -11.7, -12.7, and -9.6 rad/s for X-, Y- and Z-axes, respectively. This was distinctly different from the pattern of directional angular velocity peaks noted from the other sixteen helmets, in which primary rotation motion was noted within the coronal plane (e.g., -24.4 rad/s along the X-axis, -6.9 rad/s along the Y-axis, and -4.7 rad/s along the Z-axis for Helmet P). We speculated that the peculiar loading profile measured from Helmet Q in XRot might be related to the geometry or design of this specific helmet (e.g., this helmet was tested with an additional cover). We acknowledged the exact reason responsible for the discordant results remained unknown. To facilitate a direct comparison with our early study by Fahlstedt, et al. [39], we decided to retain Helmet Q in the current study.

In our early study by Fahlstedt, et al. [39], eight injury metrics derived from eight independently developed FE head models were used to evaluate helmet performance. It should be noted that Fahlstedt, et al. [39] studied the influence of injury metrics on helmet ranking with compounding effects from other variables among the eight FE models (e.g., material properties, element size and formulation, etc). When limiting to the few trials that ranked the helmet based on different metrics extracted from the same FE head models (i.e., MPS and CSDM extracted from the GHBMC and SIMon head models, strain and strain rate extracted from the IC head models, see the Table 2), Fahlstedt, et al. [39] reported that influence of injury metrics has a small influence on the helmet ranking outcome (Kendall’s tau: 0.84 ∼ 0.98). The current study expanded the effort by Fahlstedt, et al. [39] by involving MPS and four new tract-related strains and found that the choice of injury metrics altered the helmet ranking (Kendall’s tau: 0.42 ∼ 0.93). As no consensus has been reached on the best brain injury metric and discordant responses were noted among different FE head models, these two successive studies collectively highlighted that caution should be exercised when using computational models to rank helmet performance.

**Table. 2.** Ranking of 17 helmets using 8 different brain injury models (adapted from the study by Fahlstedt, et al. [39]).

| Model | Metric | Low-ranked ◀▶ High-ranked |  |  |  |  |  |  |  |  |  |  |  |  |  |  |  |  |
| --- | --- | --- | --- | --- | --- | --- | --- | --- | --- | --- | --- | --- | --- | --- | --- | --- | --- | --- |
| GHBMC | MPS | P | K | G | N | B | C | A | I | F | E | J | L | O | H | M | D | Q |
|  | CSDM | P | G | K | B | N | A | C | I | F | J | E | L | O | H | M | D | Q |
| IC | Strain | G | I | K | B | P | A | E | F | C | H | N | J | L | M | Q | O | D |
|  | Strain rate | G | P | K | B | N | I | C | F | A | E | J | H | L | O | M | D | Q |
| KTH | MPS | P | G | K | B | N | C | F | I | A | E | J | O | L | H | M | Q | D |
| Piper | MPS | P | G | N | K | B | C | A | F | I | E | J | L | O | H | M | D | Q |
| SIMon | MPS | P | K | B | G | N | C | A | F | E | I | J | L | H | O | M | D | Q |
|  | CSDM | P | K | N | G | B | C | A | F | I | E | J | L | O | H | M | D | Q |
| THUMS | MPS | P | K | N | B | G | A | C | I | F | J | E | L | O | H | M | D | Q |
| UCDBTM | MPS | G | B | I | P | D | K | E | F | C | H | N | A | M | J | Q | L | O |
| WHIM | MPS | P | G | K | B | N | C | A | F | I | E | J | L | O | H | M | D | Q |

Although the current study yielded some new insight into the brain strain dynamics in helmeted impacts, several caveats should be noted. First, we only involved seventeen bicycle helmets, representing a relatively small sample size than some other studies, e.g., 27 bicycle helmets by Abayazid, et al. [35], 30 bicycle helmets by Bland, et al. [15], 35 bicycle helmets by Deck, et al. [16]. The current study could be extended by involving more bicycle helmets and even other helmet types, e.g., motorcycle helmets [96], ice hockey helmets [97], and football helmets [98], etc. Second, the responses of the ADAPT model have been previously evaluated by experimental data of brain MPS and maximum principal strain rate, brain-skull relative motion, and intracranial pressure [65, 66] and showed good correlation with the brain injury pattern in a skiing accident (based on MPS) [99] and a concussive impact in American football (based on MTON) [80]. In the current study, we used the ADAPT model to predict brain responses in helmeted impacts based on MPS and four other tract-related strains. As aforementioned, only one *in vivo* imaging study [62] calculated all four types of tract-related strains with impact severity far from the loading regime in our work. Thereby, the accuracy of FE-derived tract-related strains remained to be further evaluated by their experimental counterparts measured from impact severity close to injury. The injury predictability of tract-related strains, especially the newly used MTPN, MTOS, and MTPS, needed to be further evaluated. However, it should be noted that this limitation was not necessarily specific to our study and instead represented significant challenges in the research field of brain injury biomechanics. Despite accounting for the spatial distribution of strain-based metrics in helmet evaluations, the current study neglected the temporal responses of brain deformation, as the time-accumulated strain peaks were used. Recently, Ji, et al. [79] proposed to a novel method to capture the main features of brain temporal responses by fitting the time-history brain strain curve into the Gaussian function. This method could be combined with the current study to evaluate the helmet performance with the brain dynamics in spatial and time domains collectively used. Lastly, utilizing FE simulation for helmet evaluations was often associated with enormous demands in time and resource (e.g., 4 hours when using the ADAPT head model to simulate an impact of 30 ms, with the need of 128 central processing units in a supercomputing platform). Encourage progresses have been noted on leveraging the machine learning method to instantly predict time-accumulated brain strain peaks at the regional [100] and elementwise level [101-103], and even spatially and temporally varying brain responses [104]. Of particular relevance, Ghazi, et al. [105] trained a convolution neural network (CNN) by learning elementwise brain strain under helmeted impacts simulated by the Worcester Head Injury Model (WHIM) FE head model [92]. This CNN tool was utilized to assess the performance of 23 football helmets within a fraction of second using a low-end computer [105]. Future work could consider integrating such highly efficient tools to accelerate the helmet assessment.

Several limitations in the helmet tests, that were performed early [17], should also be clarified. Firstly, the Hybrid III headform in bare condition was used to record the impact kinematics. Compared to the human head, this headfrom has unrealistic moments of inertia (MoIs) and coefficient of friction (CoF) [106-110], both of which could influence the impact kinematics and hence tissue-based strain results. For example, based on the head scans of 56 living adults, Connor, et al. [111] measured the MoIs (mean ± standard deviation) along the X (posterior-anterior), Y (left-right), and Z (inferior-superior) axes to be 189 ± 36, 200 ± 37, and 130 ± 25 kg/cm^2^, respectively, differing from those measured from the Hybrid III headform (i.e., Ixx: 161 kg/cm^2^, Iyy: 221 kg/cm^2^, Izz: 179 kg/cm^2^) [106, 108, 112]. Trotta, et al. [110] reported the CoF between the human scalp and helmet linear to be 0.29 ± 0.07, much smaller than that between the Hybrid III headform and helmet linear (0.75 ± 0.06). These limitations could be partially addressed by covering the Hybrid III headform with low-frictional materials, such as nylon stockings (CoF: 0.26 ± 0.1) [113]. Recently, a new headform (i.e., EN 17950) was developed with more biofidelic MoIs and CoF (i.e, Ixx: 196 kg/cm^2^, Iyy: 232 kg/cm^2^, Izz: 151 kg/cm^2^, CoF between the headform and liner: 0.18) [106]. This new headform is suggested to be used in future European helmet test standards, contributing to better representation of human head. Secondly, the helmet experiments were performed based on the assumption that the influence of the neck and neck and body on brain tissue responses was negligible [40, 114], as an isolated headform was used. However, the validity of this assumption was a matter of debate (see the reviews by Emsley, et al. [7] and Whyte, et al. [8]). In relation to this, the helmet’s velocity did not reach zero within 30 ms (see Fig A1 in Appendix), substantiating the absence of the deceleration phase. Several investigations [115-117] highlighted the addition of deceleration phases affected the strain response. Some studies (e.g., [105, 118, 119]) coupled the Hybrid III dummy neck with the headform in helmet testing, through which both the acceleration and deceleration phases were laboratorially presented. Critically, this neck surrogate was well known to have biofidelity issues (e.g., overly stiff in lateral bending) [120]. Chung, et al. [121] investigated two phases of the loading in an oblique helmet impact with the Hybrid III neckform and concluded the second phase (extension) was not realistic. There is ongoing work to develop more biofidelic neckforms [122, 123].

## Conclusion

This study investigated brain strain distributions and peaks in fifty-one helmeted impacts, encompassing seventeen commonly used bicycle helmets and three critical impact locations. We found that the strain distribution depended on the impact location but not on the helmet model, while the strain peak was significantly influenced by both. The helmet ranking outcome based on the strain peaks was affected by the choice of injury metrics. It was also shown that angular motion-related metrics significantly correlated with tract-related strains in the WM and MPS in the brain. Our study provided new insights into the computational brain biomechanics and helmet evaluation using tissue-based strain peaks. Our results hinted that the protective performance of helmets could be augmented by mitigating the strain peak and optimizing the strain distribution by accounting the selective vulnerability of brain subregions, although this implication required the prerequisite of region-specific injury criteria.

## Acknowledgments

This research has received funding from KTH Royal Institute of Technology (Stockholm, Sweden), the Swedish Governmental Agency for Innovation systems (VINNOVA) (no. 2023-00753), and private funds from Mips AB. The content of this article is solely the responsibility of the authors and does not necessarily represent the official views of either funding agencies. The authors also acknowledge Folksam, Sweden for sharing the bicycle helmet test results. The computational simulations were enabled by resources provided by the National Academic Infrastructure for Supercomputing in Sweden (NAISS) at the center for High Performance Computing (PDC) partially funded by the Swedish Research Council through grant agreement no. 2022-06725.

## Appendix

**Fig. A1.**
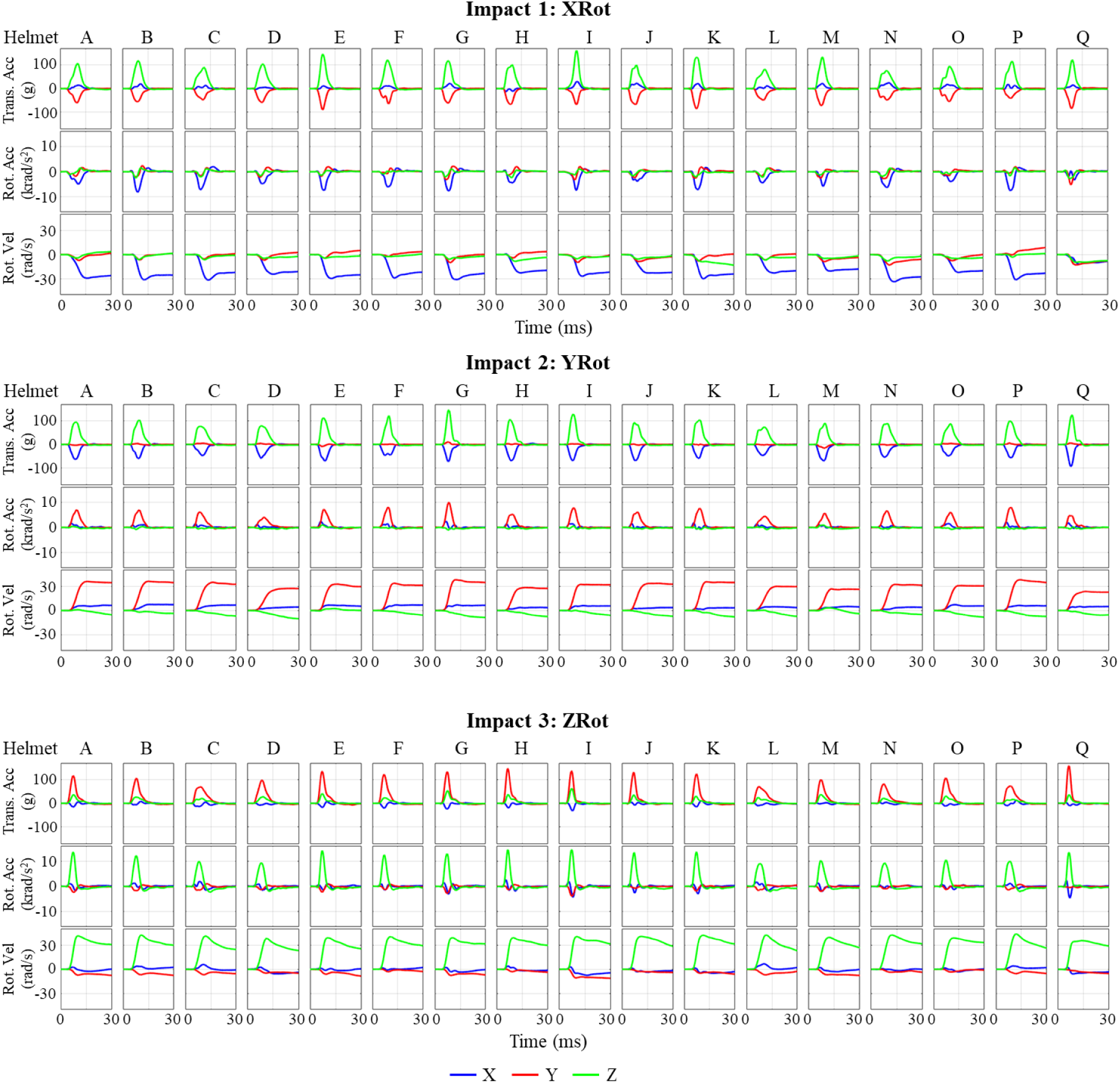
Experimental kinematics under impact 1 (i.e., XRot), impact 2 (i.e., YRot), and impact 3 (i.e., ZRot) measured from 17 helmets. The anatomical coordinate systems (i.e., XYZ in the legend) that experimental kinematics expressed with are the one in Fig 1B.

**Fig A2.**
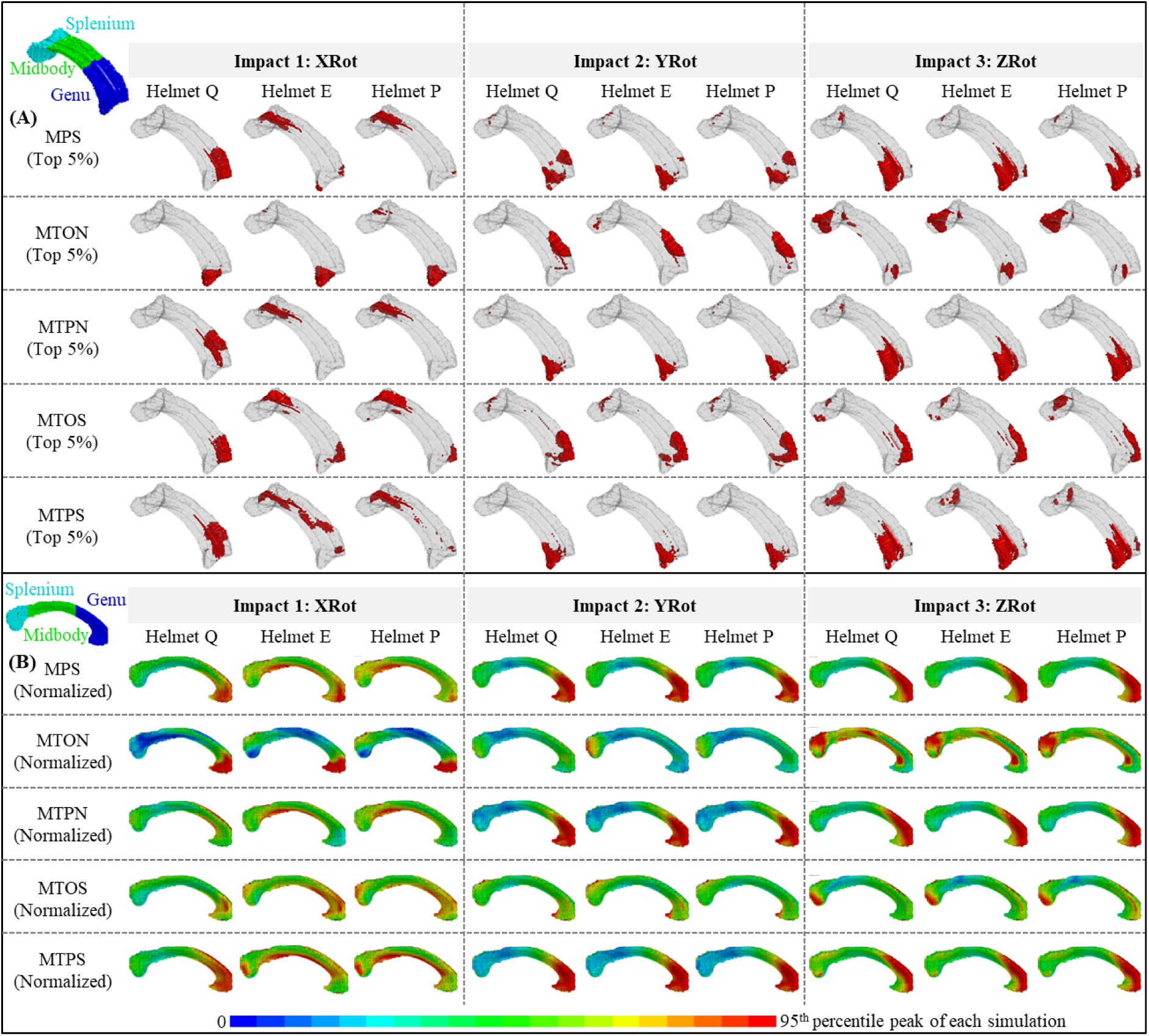
Strain distribution in the corpus callosum in nine representative impacts (i.e., Helmets Q, E, and P in XRot, YRot, and ZRot). (**A**) Elements with the MPS values and four types of tract-related strain values within the top 5%. All the figures were shown in the isometric view with an anatomical sketch (showing the splenium, midbody and genu of the corpus callosum) in the top-left corner. (**B**) Normalized strain contours. Note that, as indicated by the upper limit of the legend bar, the strain in each contour was normalized by the 95^th^ percentile strain values in the corpus callosum from the same simulation. Similarly, all the figures were shown in the side view with an anatomical sketch in the top-left corner.

**Table A1.** Values of five strain-based metrics in 51 helmeted impacts, covering 17 helmet models (Helmets A-Q) and three impact locations (XRot, YRot, and ZRot).

| (A) | Impact 1: XRot |  |  |  |  |  |  |  |  |  |  |  |  |  |  |  |  |
| --- | --- | --- | --- | --- | --- | --- | --- | --- | --- | --- | --- | --- | --- | --- | --- | --- | --- |
|  | A | B | C | D | E | F | G | H | I | J | K | L | M | N | O | P | Q |
| MPS | 0.243 | 0.273 | 0.277 | 0.206 | 0.257 | 0.248 | 0.276 | 0.207 | 0.239 | 0.198 | 0.275 | 0.197 | 0.191 | 0.306 | 0.195 | 0.270 | 0.180 |
| MTON | 0.201 | 0.225 | 0.230 | 0.168 | 0.203 | 0.198 | 0.229 | 0.162 | 0.201 | 0.166 | 0.219 | 0.163 | 0.159 | 0.259 | 0.162 | 0.207 | 0.124 |
| MTPN | 0.204 | 0.225 | 0.225 | 0.172 | 0.216 | 0.205 | 0.225 | 0.170 | 0.194 | 0.162 | 0.218 | 0.163 | 0.154 | 0.245 | 0.158 | 0.226 | 0.152 |
| MTOS | 0.176 | 0.190 | 0.196 | 0.152 | 0.178 | 0.174 | 0.192 | 0.152 | 0.172 | 0.148 | 0.189 | 0.147 | 0.141 | 0.214 | 0.147 | 0.180 | 0.124 |
| MTPS | 0.125 | 0.137 | 0.134 | 0.107 | 0.138 | 0.131 | 0.134 | 0.109 | 0.112 | 0.093 | 0.131 | 0.099 | 0.088 | 0.137 | 0.091 | 0.149 | 0.099 |
| (B) | Impact 2: YRot |  |  |  |  |  |  |  |  |  |  |  |  |  |  |  |  |
|  | A | B | C | D | E | F | G | H | I | J | K | L | M | N | O | P | Q |
| MPS | 0.323 | 0.328 | 0.305 | 0.224 | 0.301 | 0.321 | 0.365 | 0.254 | 0.308 | 0.297 | 0.333 | 0.257 | 0.248 | 0.300 | 0.288 | 0.354 | 0.205 |
| MTON | 0.159 | 0.166 | 0.153 | 0.112 | 0.152 | 0.159 | 0.181 | 0.123 | 0.152 | 0.142 | 0.157 | 0.127 | 0.118 | 0.144 | 0.147 | 0.176 | 0.103 |
| **MTPN** | 0.293 | 0.298 | 0.280 | 0.211 | 0.269 | 0.287 | 0.327 | 0.233 | 0.276 | 0.271 | 0.299 | 0.239 | 0.221 | 0.273 | 0.264 | 0.320 | 0.190 |
| **MTOS** | 0.176 | 0.180 | 0.169 | 0.128 | 0.166 | 0.176 | 0.197 | 0.142 | 0.168 | 0.162 | 0.178 | 0.144 | 0.136 | 0.163 | 0.163 | 0.193 | 0.119 |
| **MTPS** | 0.212 | 0.212 | 0.201 | 0.156 | 0.196 | 0.205 | 0.227 | 0.173 | 0.198 | 0.199 | 0.218 | 0.176 | 0.169 | 0.200 | 0.189 | 0.226 | 0.139 |
| **(C)** | **Impact 3: ZRot** |  |  |  |  |  |  |  |  |  |  |  |  |  |  |  |  |
| **Helmets** | **A** | **B** | **C** | **D** | **E** | **F** | **G** | **H** | **I** | **J** | **K** | **L** | **M** | **N** | **O** | **P** | **Q** |
| **MPS** | 0.427 | 0.431 | 0.417 | 0.393 | 0.406 | 0.410 | 0.410 | 0.403 | 0.430 | 0.415 | 0.430 | 0.426 | 0.404 | 0.423 | 0.395 | 0.435 | 0.370 |
| **MTON** | 0.314 | 0.321 | 0.313 | 0.284 | 0.295 | 0.301 | 0.297 | 0.298 | 0.317 | 0.303 | 0.313 | 0.321 | 0.297 | 0.310 | 0.289 | 0.325 | 0.264 |
| **MTPN** | 0.361 | 0.367 | 0.356 | 0.323 | 0.341 | 0.347 | 0.341 | 0.339 | 0.359 | 0.353 | 0.364 | 0.366 | 0.340 | 0.361 | 0.334 | 0.374 | 0.307 |
| **MTOS** | 0.267 | 0.272 | 0.267 | 0.245 | 0.256 | 0.259 | 0.255 | 0.257 | 0.264 | 0.263 | 0.269 | 0.272 | 0.256 | 0.268 | 0.253 | 0.277 | 0.234 |
| **MTPS** | 0.220 | 0.222 | 0.215 | 0.202 | 0.209 | 0.214 | 0.210 | 0.208 | 0.220 | 0.219 | 0.225 | 0.220 | 0.210 | 0.223 | 0.207 | 0.225 | 0.195 |

